# Neofunctionalisation of basic helix loop helix proteins occurred when plants colonised the land

**DOI:** 10.1101/514109

**Authors:** Clémence Bonnot, Alexander J. Hetherington, Clément Champion, Holger Breuninger, Steven Kelly, Liam Dolan

**Affiliations:** Department of Plant Sciences, University of Oxford, South Parks Road, Oxford, OX1 3RB, UK

**Keywords:** basic helix loop helix (bHLH), *Chara braunii*, *Coleochate nitellarum*, Land plants, *ROOT HAIR DEFECTIVE SIX-LIKE* (RSL), Filamentous rooting cells, Streptophyte algae, Neofunctionalisation

## Abstract

*ROOT HAIR DEFECTIVE SIX-LIKE (RSL)* genes control the development of structures – rhizoids, root hairs, gemmae, mucilage papillae – that develop from single cells at the surface of diverse groups of land plants. RSL proteins constitute a subclass (VIIIc) of the basic helix loop helix (bHLH) class VIII transcription factor family. We set out to determine if the function of RSL genes in the control of cell differentiation in land plants was inherited from streptophyte algal ancestor. The Charophyceae are a monophyletic class of streptophyte algae with tissue-like structures and rhizoids. We identified the single class VIII bHLH gene from the charophyceaen alga *Chara braunii* (Cb*bHLHVIII*). Phylogenetic analysis suggests that this protein is sister to the RSL (bHLH subclass VIIIc) proteins and together they constitute a monophyletic group. Expression of Cb*bHLHVIII* does not compensate for loss of the RSL function in either Marchantia polymorpha or Arabidopsis thaliana. Furthermore, CbbHLHVIII is expressed at sites of morphogenesis in *C. braunii* – the apices, nodes and gametangia – but not in rhizoids. This indicates that *C. braunii* class VIII protein is functionally different from land plant RSL proteins; they control rhizoid development in land plants but not in the charophycean algae. These data are consistent with the hypothesis that RSL proteins and their function in the differentiation of cells at the plant surface evolved in the lineage leading to land plants after the divergence of the land plants and *C. braunii* from their last common ancestor. This may have occurred by neofunctionalisation at or before the colonisation of the land by streptophytes.

## INTRODUCTION

The colonisation of the land by streptophytes and the subsequent radiation of morphological diversity among the land plants was a major transition in Earth history (Kenrick & Crane, 1997; Pires & Dolan, 2012). The simple body plans of extant streptophyte algae has led to the hypothesis that the body plan of the common ancestor of the land plants was simple. It is hypothesized that plants then underwent morphological diversification during and after the colonisation of the dry continental surfaces (Graham *et al.*, 2000; Delaux *et al.*, 2012; Harrison, 2017). These adaptions to life on the land contributed to the establishment of the first complex terrestrial ecosystems by 407 Ma and led to the terrestrial ecosystems that exist today (Gibling & Davies, 2012; Lenton *et al.*, 2012).

The sequencing of land plant (Rensing *et al.*, 2008; Banks *et al.*, 2011; Bowman *et al.*, 2017) and streptophyte algae genomes (Hori *et al.*, 2014; Ju *et al.*, 2015; Nishiyama *et al.*, 2018) has led to the hypothesis that the evolution of the morphological diversity resulted from an increase in the number of regulatory genes in gene families (Floyd & Bowman, 2007; Lang *et al.*, 2010; Pires & Dolan, 2010a,b; Bowman *et al.*, 2017; Lehti-Shiu *et al.*, 2017). For example, transcription factor family number is higher in land plants than in streptophyte algae (Tanabe *et al.*, 2005; Navaud *et al.*, 2007; Chanderbali *et al.*, 2015; Catarino *et al.*, 2016; Nishiyama *et al.*, 2018). It is also likely that the function of regulatory genes will have changed during the course of the transition to land. Such changes could result from genes assuming new functions (neofunctionalisation) or dividing their functions among their descendants (sub-functionalisation) (Prince & Pickett, 2002; Rensing, 2014).

The development of a diversity of morphological structures that form from single cells in the surface cell layer of organs is regulated by RSL class 1 genes (also known as subclass VIIIc1 basic helix loop helix) in a diversity of land plants (Honkanen & Dolan, 2016). For example, RSL class 1 genes positively regulate the development of rhizoids, mucilage papillae and gemmae (asexual propagules) in the liverwort *Marchantia polymorpha* (Proust *et al.*, 2016). Orthologs positively regulate the development of rhizoids and mucilage papillae in the moss *Physcomitrella patens* (Menand *et al.*, 2007; Jang *et al.*, 2011; Proust *et al.*, 2016). Class 1 RSL genes positively regulate the development of root hairs in diverse groups of angiosperms including *Arabidopsis thaliana* (Masucci & Schiefelbein, 1994; Menand *et al.*, 2007) and the grasses *Oryza sativa* and *Brachypodium distachyon* (Kim & Dolan, 2016; Kim *et al.*, 2017). Expression of RSL class 1 genes from one taxa of land plant can compensate for the loss of function in another taxa (Menand *et al.*, 2007; Kim & Dolan, 2016; Proust *et al.*, 2016; Kim *et al.*, 2017). For example, expression of *M. polymorpha* Mp*RSL1* gene using the cauliflower mosaic virus 35S promoter (35S) in the root hairless At*rhd6* At*rsl1* mutants of A. thaliana restores root hair development (Proust *et al.*, 2016). This demonstrates that not only do RSL class 1 genes control the development of these structures in diverse land plants, but also that the function of the proteins has been conserved during the course of land plant evolution.

RSL class 1 genes (subclass VIIIc1) are members of class VIII basic helix loop helix (bHLH) transcription factors (Menand *et al.*, 2007; Pires & Dolan, 2010a; Proust *et al.*, 2016). Class VIII bHLH proteins includes two other subclasses, subclass VIIIa and subclass VIIIb (Heim *et al.*, 2003; Pires & Dolan, 2010a; Catarino *et al.*, 2016). The functions of subclass VIIIa proteins are unknown (Pires & Dolan, 2010a). Subclass VIIIb includes the HECATE-related transcription factors that control fruit development in *A. thaliana* (Liljegren *et al.*, 2004; Gremski *et al.*, 2007; Kay *et al.*, 2013). Together the subclasses VIIIa, VIIIb and VIIIc constitute a monophyletic group of proteins that is conserved among land plants. While the function of RSL class 1 genes (subclass VIIIc) have been shown to be conserved among diverse groups of land plants (Menand *et al.*, 2007; Jang *et al.*, 2011; Kim & Dolan, 2016, p. 20; Proust *et al.*, 2016; Kim *et al.*, 2017), the functions of subclass VIIIb have only been defined in *A. thaliana* to date (Liljegren *et al.*, 2004; Gremski *et al.*, 2007; Kay *et al.*, 2013).

Since RSL class 1 genes control the development of rhizoids and root hairs in the mosses, liverworts and angiosperms, it is hypothesized that RSL class 1 genes control the development of the rhizoidal rooting structure in the common ancestor of the land plants (Menand *et al.*, 2007; Proust *et al.*, 2016). This was one of a suite of new functions that evolved early in land plant evolution, which was a key adaptation to life on the continental surfaces of the planet (Kenrick & Crane, 1997; Delaux *et al.*, 2012). To investigate the origin of this regulatory mechanism we searched the genome (Nishiyama *et al.*, 2018) and transcriptomes of the streptophyte alga *Chara braunii* for genes encoding class VIII bHLH proteins. *C. braunii* is a member of the Charophyceae, the only class of streptophyte algae in which tissue-like structures and rhizoids develop (Smith & Allen, 1955; Pickett-Heaps, 1975; Graham & Wilcox, 2000; Nishiyama *et al.*, 2018). We identified a single class VIII gene (Cb*bHLHVIII*) that is sister to the land plant RSL (subclass VIIIc) transcription factors. The expression of Cb*bHLHVIII* in *M. polymorpha* or *A. thaliana* mutants does not compensate for the loss of endogenous RSL function, indicating that *C. braunii* class VIII protein is functionally different from the land plant RSL proteins. Furthermore, Cb*bHLHVIII* is expressed in regions of organogenesis - apices and young nodes of the thallus, and gametangia - but not in rhizoids or in rhizoids morphogenesis zones. This is consistent with the hypothesis that Cb*bHLHVIII* regulates development at *C. braunii* morphogenetic centres, but does not control rhizoid differentiation. We conclude that class VIII proteins evolved the ability to control the differentiation of cells such as rhizoids in the land plant lineage after the divergence of *C. braunii* and land plants from their last common ancestor. This is consistent with a model in which neofunctionalisation of class VIII proteins occurred during the increase in morphological diversity that occurred during the transition to land.

## MATERIALS AND METHODS

### Sequence identification

To identify class VIII sequences in *Chara braunii* and *Coleochaete nitellarum*, we used the and plants bHLH class VIII protein sequences of *Arabidopsis thaliana*, *Selaginella moelendorfii*, *Physcomitrella patens* and *Marchantia polymorpha* published previously (Catarino *et al.*, 2016) as queries in TBlastN searches (Altschul *et al.*, 1990) of *Coleochaete nitellarum* transcriptome (Bonnot *et al.*, 2017) and *C. braunii* genome (Nishiyama *et al.*, 2018) and transcriptome (see “Construction of the *C. braunii* transcriptome” hereafter). No E-value thresholds were used. All hits were manually investigated. Transcripts and genomic sequences were aligned. Amino acid sequences were predicted from transcripts using ExPASy translate (ExPASy, University of Geneva, Geneva, Switzerland) and analysed with SMART (http://smart.embl-heidelberg.de) and Pfam (http://pfam.sanger.ac.uk/search) to determine if a bHLH domain was coded in each transcript. Reciprocal TBlastN searches (Altschul *et al.*, 1990) were conducted on NCBI (http://blast.ncbi.nlm.nih.gov/Blast.cgi) to verify if the hits containing a bHLH domain belonged to the bHLH class VIII family.

Unique bHLH class VIII gene candidates (later named Cn*bHLHVIII* and Cb*bHLHVIII*) were found in the *C. nitellarum* transcriptome (transcript_12969), and in the *C. braunii* transcriptome (Cb_Transcript_119934) and genome (CHBRA233g00280) respectively. We verified the sequence of Cn*bHLHVIII* and Cb*bHLHVIII* transcripts by PCR (primers listed in Table 1) using cDNA from whole plant total RNA and Sanger sequencing (Source Bioscience, Nottingham, UK). The transcript encoding CnbHLHVIII was 2769 nucleotides long and contained a 1725 nucleotide CDS coding for a 574 amino acid protein (Supplementary Data 1). The transcript encoding CbbHLHVIII was 6103 nucleotides long and contained a 3468 nucleotides CDS coding for a 1155 amino acids protein (Supplementary Data 1). The CDS and protein sequences of CnbHLHVIII and CbbHLHVIII were uploaded on Genbank under the accession numbers MK292332 and MK292331 respectively.

**Table 1.**
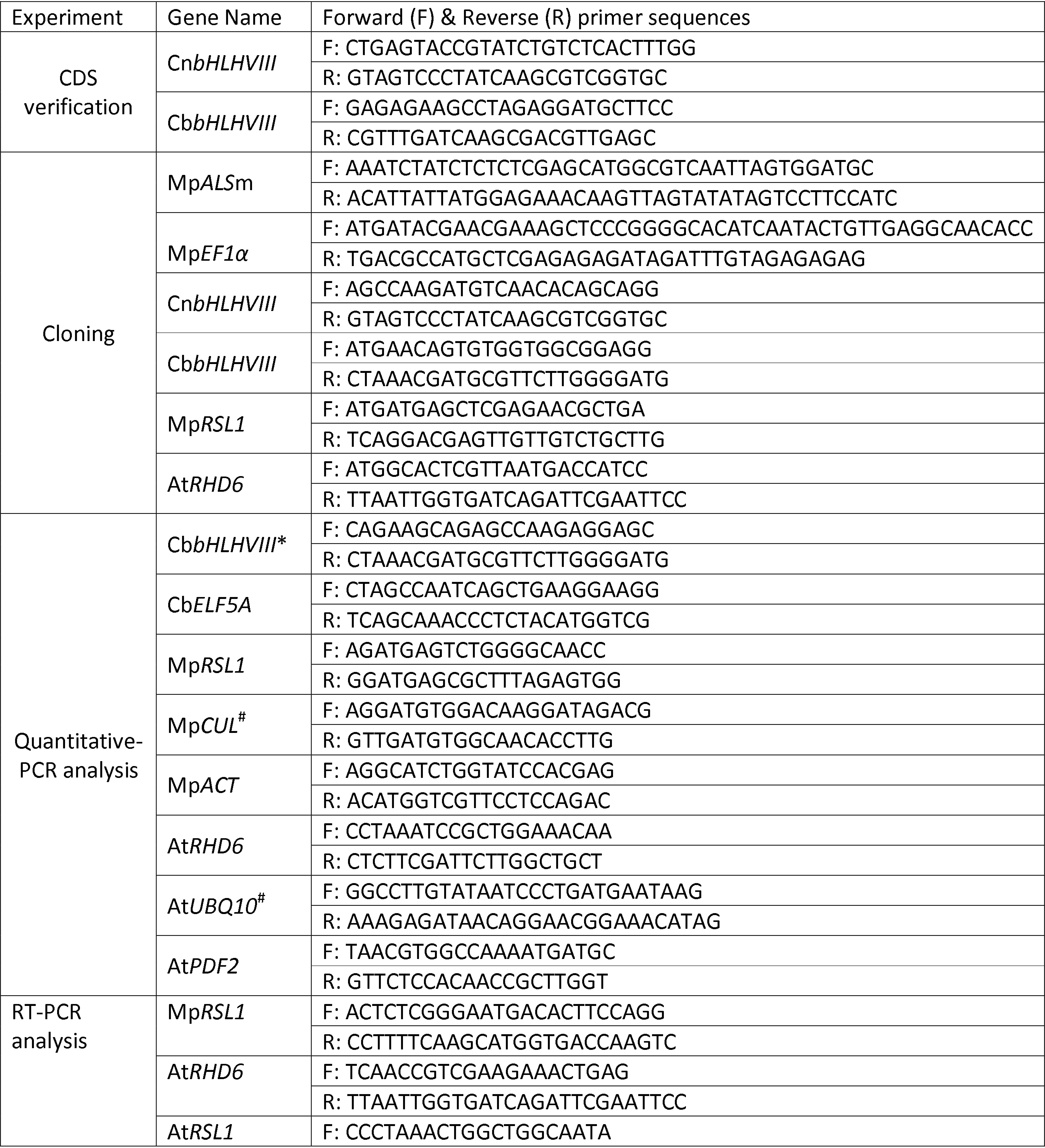
List of primers for cloning, vector construction and expression analysis.

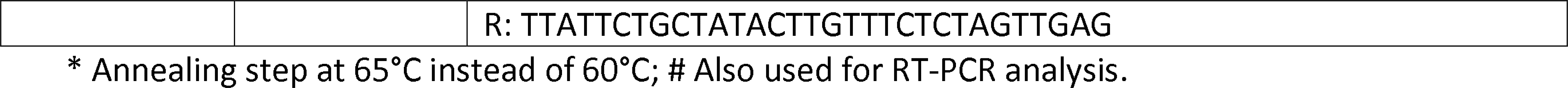

### Phylogeny and protein domain analysis

We used the iterative refinement method L-INS-i (Katoh *et al.*, 2005) on MAFFT--add (Katoh & Frith, 2012; Katoh & Standley, 2013) to add the full length protein sequences of CnbHLHVIII and CbbHLHVIII to the previously published alignment of full length bHLH proteins from the land plants *A. thaliana*, *Oryza sativa*, *S. moelendorfii*, *P. patens* and *M. polymorpha*, the streptophyte alga *Klebsormidum flaccidum*, the chlorophyte algae *Volvox carteri*, *Chlamydomonas reinhardtii*, *Chlorella variabilis* and *Ostreococcus tauri* and, the rodophyte alga *Cyanidioschyzon merolae* (Supplementary Data 2) (Catarino *et al.*, 2016). The alignment was trimmed manually with BioEdit v.7.1 (Hall, 1999) to remove the non-conserved regions (Supplementary Data 3). Maximum-likelihood analysis was carried-out with PhyML 3.0 (Guindon *et al.*, 2010), with the Jones Taylor and Thornton (JTT) amino acid substitution model (Jones *et al.*, 1992), on the complete set of aligned and trimmed proteins (Archaeplastida bHLH phylogeny; Figure 1.A and Supplementary Figure S1) and on a reduced set (Supplementary Data 4) containing only the members of the bHLH class VIII family, CnbHLHVIII and CbbHLHVIII (bHLH class VIII phylogeny; Figure 1.B and Supplementary Figure S2). The statistical branch support values were calculated with an approximate likelihood ratio test (aLRT) (Guindon *et al.*, 2010). Trees were visualised using FigTree (http://tree.bio.ed.ac.uk/software/figtree/). The presence of the bHLH class VIII conserved motifs was assessed using the amino acid alignment (Supplementary Figure S3) and MEME (Multiple Em for Motif Elicitation) (Bailey & Elkan, 1994) with a constraint of 1 to 4 conserved motifs on members of each subclasses (VIIIa, HECATE, RSL class 1, RSL class 2 and RSL class 3) associated with CnbHLHVIII and CbbHLHVIII. The LOGO representation of the amino-acid sequence of each bHLH class VIII conserved motif was produced using WebLogo (https://weblogo.berkeley.edu).

**Figure 1.**
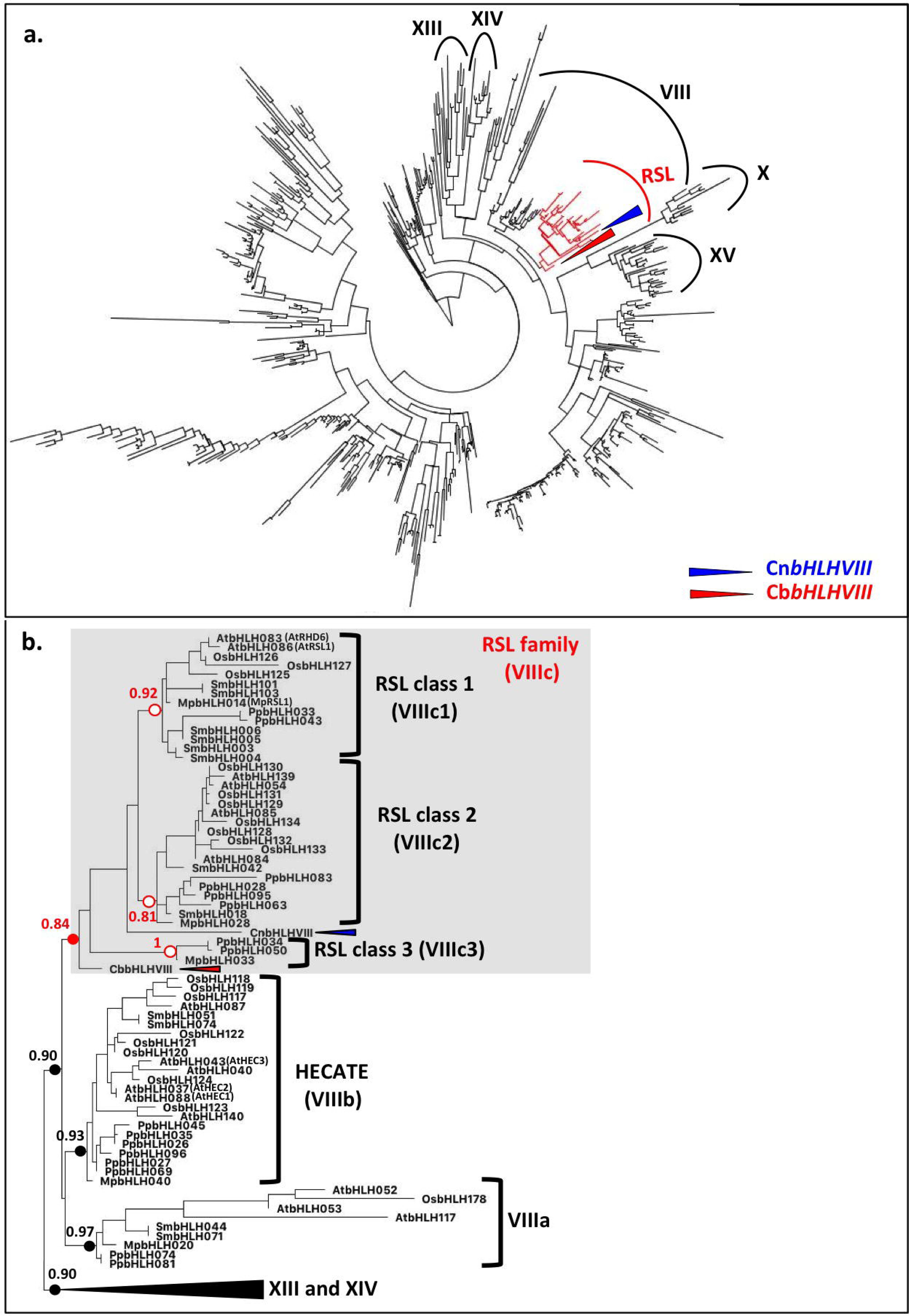
CbbHLHVIII is sister to land plant class VIII bHLH transcription factors. **a.** Unrooted maximum-likelihood tree of the Archaeplastida bHLH transcription factors. The, the complete class VIII bHLH family including the RSL (subclass VIIIc; in red) and related bHLH families (class X, XV, XIII and XIV) are marked with ellipses. CbbHLHVIII and CnbHLHVIII are marked with red and a blue triangle respectively. **b.** Maximum-likelihood tree of the class VIII bHLH transcription factors. Tree rooted with the bHLH class XIII and XIV sequences. The approximate likelihood ratio test (aLRT) support values are given for the major nodes of the tree and marked by a red circle. The red and the blue triangles mark Cb*bHLHVIII* and CnbHLHVIII respectively. *A. thaliana* (At), *O. sativa* (Os), *S. moelendorfii* (Sm), *P. patens* (Pp) and *M. polymorpha* (Mp).

### Plant growth conditions

*C. nitellarum* (strain provided by the Skidmore Algal collection of Professor David Domozych) were grown in cell culture flasks (cell star, 250 mL, filter cap; Greiner Bio-One, Kremsmunster, Austria) in 75 mL of liquid NaNO_3_-Bold Basal Medium (Nichols, 1973) supplemented with 1g.L-1 of NaNO_3_, renewed every six weeks, under a cycle of 8h:16h 1 dark:light (38 μmol.m^−2^.s^−1^) at 23°C at 45 rpm.

*C. braunii* (strain S276) (Nishiyama *et al.*, 2018) was grown under axenic conditions on a metallic net immersed in modified Forsberg liquid medium (0.56 mg.L^-1^ Na_2_HPO_4_, 0.112 g.L^−1^ Ca(NO_3_)_2_.3H_2_O, 0.1 g.L^−1^ MgSO_4_.7H_2_O, 0.0174 g.L^−1^ Na_2_SiO_3_.5H_2_O, 0.03 g.L^−1^ KCl, 0.02 g.L^−1^ Na_2_CO_3_, 2 ug.L^−1^ MnCl_2_, 2 ug.L^−1^ CoCl_2_, 4 ug.L^−1^ CuCl_2_, 0,4 mg.L^−1^ FeCl_2_, 0,1 mg.L^−1^ ZnCl_2_, 0,1 mg.L^−1^ NaMoO_4_, 0.4 mg.L^−1^ H_3_BO_3_, 0.5 g.L^−1^ TRIS, 0,02 mg.L^−1^ Nitrilo triacetic acid (NTA)) (Forsberg, 1965) pH 7.8 supplemented with Kao and Michayluk vitamin solution (100X) (#K3129; Sigma, St-Louis, MO, USA) at 23°C under a light cycle of 8 h : 16 h, dark : light (38 μmol.m^−2^.s^−1^).

*M. polymorpha* wild type accessions Takaragaike-1 (Tak-1) male and Takaragaike-2 (Tak-2) female (Ishizaki *et al.*, 2008) and the loss of function Mp*rsl1-1* mutant (Proust *et al.*, 2016) were used in this study. Meristem-containing thallus fragments of axenic Mp*rsl1-1* and wild type and, surface sterile spores produced from the cross of Tak-1 and Tak-2 were transformed with binary vectors (see vectors construction hereafter) using agrobacterium (strain GV3101) following published co-cultivation protocols (Ishizaki *et al.*, 2008; Kubota *et al.*, 2013). After transformation, regenerated thalli or sporelings were selected on ½ Johnson’s medium 1% agar supplemented with 100 μg.mL^−1^ of cefotaxime and 10 μg.mL^−1^ of hygromycin or 10 μM of chlorsulfuron depending on the vector (see vector construction hereafter). For phenotypic and expression analyses, Tak-1, Tak-2 and Mprsl1-1 thalli, Tak-1 and Tak-2 sporelings, transformed sporelings or thalli were grown in axenic conditions on ½ Johnson’s medium 0.7% agar at 23°C under continuous illumination (38 μmol.m^−2^.s^−1^).

Wild type *A. thaliana* Columbia (Col-0) and the loss of function At*rhd6-3* At*rsl1-1* double mutant (Menand *et al.*, 2007) were used in this study. Plants were transformed by floral deep (Zhang *et al.*, 2006) with binary vectors (see vectors construction hereafter) using agrobacterium (strain GV3101) cultures. Before, in vitro culture seeds were surface sterilized ten minutes with a solution of 70% Ethanol and 0.1% Triton X-100 and ten minutes with a solution of 99% ethanol. After transformation seeds were selected on MS medium 1% agar supplemented with 50 ug.mL^−1^ of hygromycin. For phenotypic and expression analyses, seeds of wild type, mutants and of three T2 transformant lines for each construct were grown as previously described (Breuninger *et al.*, 2016).

### Phenotypic analysis and image acquisition

Transmitted light microscopy images of *C. braunii*, *M. polymorpha* and *A. thaliana* were captured using a camera (Leica DFC310 FX; Leica, Wetzlar, Germany) mounted on a dissecting microscope (Leica M165 FC; Leica, Wetzlar, Germany). For *A. thaliana*, 15 plants per line were phenotyped 10 days after germination. For each line of *M. polymorpha* over-expressing 35S:At*RHD6*, Mp*RSL1* and Cb*bHLHVIII* in the wild type background, 5 plants were phenotyped four weeks after spore transformation. 20 plants were phenotyped for each line generated by transforming the vector into the *M. polymorpha* Mp*rsl1* knockout mutant background, seven and ten weeks after thallus regeneration.

*C. braunii* rhizoid nodal growth pattern was analysed on 107 nodes (32 N_1_, 26 N_2_, 22 N_3_, 15 N_4_ and 12 N_older_) from 13 plants. The stage (N_1_, N_2_, N_3_, N_4_ or N_older_) and the presence or absence of initiating, unicellular elongating, multicellular and branched rhizoids were noted for each node. The percentage of node of each stage (N_1_, N_2_, N_3_, N_4_ and N_older_) displaying initiating, unicellular elongating, multicellular and branched rhizoids were calculated. Most nodes displayed rhizoids at different stages and were therefore used in the calculation of the percentage of node for each of the rhizoid categories they displayed.

### RNA extraction, cDNA synthesis

Total RNA was extracted with a mir™Vana Kit (#AM1560; Thermo Fisher scientific, Waltham, MA, USA) from frozen *C. braunii* whole plants for cDNA synthesis and cloning. Total RNA for transcriptomes was isolated using the same protocol from frozen rhizoids, green thalli with attached gametangia and whole plants. mRNA was extracted with a Dynabeads^®^mRNA DIRECT™Kit (#61011; Thermo Fisher scientific, Waltham, MA, USA) from ground frozen tissues collected as follows. Two weeks after propagation, rhizoids grown below a metallic net were cut and flash frozen in liquid nitrogen for total RNA and mRNA extraction. *C. braunii* thallus cleaned from any remaining rhizoids with gametangia (for total RNA extraction) or without (for mRNA extraction) and, thallus parts (apices, nodes, branches, gametangia and zygotes were collected by hand under a dissecting microscope (Leica M165 FC; Leica, Wetzlar, Germany) and flash frozen in liquid nitrogen. Total RNA and mRNA were DNase treated using the Turbo DNase™Kit (#AM2238; Thermo Fisher scientific, Waltham, MA, USA) according manufacturer’s instructions. cDNAs were synthesized from 5 μg of total RNA or 20 ng of mRNA in 20 uL reaction volume using the SuperScript III First-strand synthesis System (#18080051; Thermo Fisher scientific, Waltham, MA, USA) with the oligo (dT_20_) provided.

Total RNA was extracted with Direct-Zol RNA Miniprep Kits (#R2060; Zymo research, Irvine, CA, USA) from ground frozen whole plant for *C. nitellarum* and *M. polymorpha* and, from roots for *A. thaliana*. *C. nitellarum*, *M. polymorpha* and *A. thaliana* samples of total RNA were DNase treated using the Turbo DNase-free™ Kit (#AM1907; Thermo Fisher scientific, Waltham, MA, USA). cDNAs were synthesised from 1 μg of total RNA in a 20 μL reaction volume using 200 U of Protoscript II reverse transcriptase (#M0368S; NEB, Ipswich, MA, USA), oligo (dT_17_) at 2 μM and the NEB Murine RNase inhibitor (#M0314; NEB, Ipswich, MA, USA).

### Construction of the *C. braunii* transcriptome

The whole plant *C. braunii* transcriptome was produced by the sequencing of ten strand-specific cDNA libraries (two whole plants libraries, two green thallus libraries and four rhizoid libraries) with an insert size of 500 bp, using a paired-end read length of 2 × 100 bp on Illumina Hiseq2000. GATC Biotech Ltd (Eurofingenomics, Konstanz, Germany) performed the preparation of the libraries from 2 μg of total RNA and sequencing.

Raw reads were quality trimmed with Trimmomatic-0.32 (Bolger *et al.*, 2014), to remove Illumina adaptors and low quality tails. Ribosomal RNA was filtered out with Sortmerna-1.9 (Kopylova *et al.*, 2012). Reads were further error corrected using Allpaths-LG-4832 (Butler *et al.*, 2008) (with setting PAIRED_SEP=−20 and ploidy = 1). Trimmed and corrected reads were normalised using Khmer-0.7.1 with a khmer size of 31. Before assembly, paired end reads were stitched together using Allpaths-LG-4832 (Butler *et al.*, 2008). A de novo transcriptome assembly was made with the cleaned, stitched reads using SGA (Simpson & Durbin, 2012), SSPACE-v3 (Boetzer *et al.*, 2011), and CAP3 (Huang & Madan, 1999). Finally assembled scaffolds were corrected using Pilon-1.6 (Walker *et al.*, 2014). The transcriptome assembly of *C. braunii* consisted of 117,611 transcripts with a mean sequence length of 749 bp. 21,917 of the transcripts were over 1kb in length. This Transcriptome Shotgun Assembly project has been deposited at DDBJ/EMBL/GenBank under the accession GGXX00000000. The version described in this paper is the first version, GGXX01000000.

The expression level of each transcript was next quantified. Raw reads were quality trimmed with Trimmomatic-0.32 (Bolger *et al.*, 2014), Sortmerna-1.9 (Kopylova *et al.*, 2012) and BAYESHAMMER (SPADES-3.5.0) (Nikolenko *et al.*, 2013). Corrected reads were mapped to the transcriptome using Salmon 0.9.1 (Patro *et al.*, 2017).

### Cloning and vector construction

The CDSs encoding CbbHLHVIII, CnbHLHVIII, MpRSL1 and AtRHD6 were amplified respectively from *C. braunii*, *C. nitellarum*, *M. polymorpha* and *A. thaliana* cDNA made from total RNA, using the high-fidelity polymerase Phusion (#F530L; Thermo Fisher Scientific, Waltham, MA, USA) with the primers listed in Table 1. For *C. braunii* and *C. nitellarum* we used a slow elongation rate of 70s.Kb^−1^. The CDSs of Cb*bHLHVIII*, Mp*RSL1* and At*RHD6* were then cloned into pCR8-GW-TOPO vectors (#K2500-20; Thermo Fisher Scientific) to establish Gateway™ (GW) entry vectors. For *A. thaliana* transformations, the pCR8 entry vectors containing the CDSs encoding CbbHLHVIII and AtRHD6 were recombined with the destination vector pCambia p35S:GW:T (Breuninger *et al.*, 2016) by LR reactions (#12538-200; Thermo Fisher Scientific). For *M. polymorpha* transformations, the entry vectors containing the CDSs encoding CbbHLHVIII and MpRSL1 were recombined by LR reactions (#12538-200; Thermo Fisher scientific, Waltham, MA, USA) with the destination vectors pCambia p35S:GW:T (carrying the resistance to hygromycin) (Breuninger *et al.*, 2016) and pCambia pMpEF1α:GW:T carrying a mutated version of the MpALSm gene driven by the *p35S* promoter and conferring plant resistance to chlorsulfuron. This vector was generated by splicing PCR products of the Mp*ALSm* gene with the 35S promoter together with the Mp*EF1α* promoter into a XhoI/PstI-digeststed pCAMBIA backbone using the Clontech In-Fusion kit (Clontech Cat. #: 638909) and the primers indicated in Table 1. Subsequently, this vector was converted into a Gateway™ destination vector using the inserted SmaI site next to the MpEF1α promoter using the Gateway™ Vector Conversion System (Thermo Fisher Cat. #: 11828029).

### Expression analysis

Quantitative PCR analyses were done on a 7300 Applied Biosystems thermocycler (Thermo Fisher scientific, Waltham, MA, USA) using the SensiFAST™ Sybr^®^ Hi-ROX Kit (#BIO-92020; Bioline, London, UK). Amplification was performed in a reaction volume of 10 μL containing 500 nM of each of the forward and reverse primers and 4 μL of a two-fold dilution of *C. braunii* cDNA synthesized from mRNA or of ten-fold dilutions of *M. polymorpha* and *A. thaliana* cDNA synthesized from total RNA. Data analysis were carried our as previously described (Saint-Marcoux *et al.*, 2015) using the reference genes At*UBQ10* and At*PDF2* (Czechowski *et al.*, 2005), Mp*ACT* and Mp*CUL* (Saint-Marcoux *et al.*, 2015) and, CbELF5a (CHBRA130g00470). All primer used are listed in Table 1.

## RESULTS

### There is a single class VIII basic helix loop helix transcription factor in the *Chara braunii* genome

To determine if genes encoding class VIII bHLH transcription factors control rhizoid development in charophyceaen algae, we searched a *C. braunii* transcriptome for similar sequences. We isolated RNA from whole *C. braunii* plants, green parts and from rhizoids. We sequenced the RNA and assembled a transcriptome with an N50 value of 1229 bp and a mean length of 749 bp (Table 2). Using the TBLASTN algorithm and the class VIII protein sequences of *A. thaliana*, *Selaginella moelendorffi*, *Physcomitrella patens* and *M. polymorpha* (Catarino *et al.*, 2016) as queries, we identified a single nucleotide sequence encoding a predicted protein of 1155 amino acids similar to the land plant class VIII proteins (Cb_Transcript_119934) (Supplementary Data 1). We further searched the *C. braunii* genome and transcriptome assemblies (Nishiyama *et al.*, 2018), using the same queries and the complete sequence of the putative *C. braunii* class VIII bHLH protein as a query but did not identify further new class VIII sequences. We also identified a putative class VIII bHLH protein from a *Coleochaete nitellarum* transcriptome (Bonnot *et al.*, 2017) (Supplementary Data 1). No putative class VIII bHLH sequence was found in *Klebsormidium flaccidum* (Hori *et al.*, 2014; Catarino *et al.*, 2016).

**Table 2.** Transcriptome parameters

|  |  |
| --- | --- |
| Number of sequences | 117611 |
| Median sequence length | 395 nucleotides |
| Mean sequence length | 749 nucleotides |
| Max sequence length | 21672 nucleotides |
| Min sequence length | 200 nucleotides |
| No. sequences > 1kb | 21917 |
| No. sequences > 10kb | 63 |
| No. gaps | 660 |
| N50 | 1229 |
| Combined sequence length | 88124568 |

To test if the putative *C. braunii* and *C. nitellarum* class VIII bHLH proteins belong to the class VIII bHLH family, we aligned the sequences around their bHLH domain with bHLH sequences from 11 Archaeplastida taxa and generated a phylogenetic tree using the maximum likelihood statistics (Figure 1 and Supplementary Figure S1 and S2) (Catarino *et al.*, 2016). The topology of the tree indicated that the *C. braunii* putative class VIII bHLH protein was sister to the RSL (subclass VIIIc) proteins while *C. nitellarum* putative class VIII bHLH protein was sister to the RSL class 1 (subclass VIIIc 1) and class 2 (subclass VIIIc 2). Since the *C. braunii* and *C. nitellarum* proteins constituted a monophyletic group with the other class VIII bHLH proteins, we conclude that the *C. braunii* and *C. nitellarum* proteins are class VIII members, and designated them respectively CbbHLHVIII and CnbHLHVIII. These data indicate that a single gene encoding a class VIII bHLH protein existed in the last common ancestor of *C. braunii*, *C. nitellarum* and the land plants. They also demonstrate that a single copy class VIII bHLH gene is present in both *C. braunii* that develops rhizoids and in *C. nitellarum* that does not develop rhizoids.

### CbbHLHVIII and CnbHLHVIII lack the conserved motifs present in the land plant class VIII proteins

Land plant class VIII bHLH proteins are characterised by distinct, conserved motifs near and in the bHLH domain (Figure 2). They possess a very conserved bHLH domain (Figure 2.A and Supplementary Figure S3) containing an atypical basic domain characterised by the presence of a conserved Alanine (A^210^ in AtRHD6) at position 9 of the bHLH domain (Figure 2.A and Supplementary Figure S3) that is not found in the other bHLH classes (Supplementary Figure S4 and Data S2) (Heim *et al.*, 2003; Liljegren *et al.*, 2004; Gremski *et al.*, 2007). There is an atypical basic domain, typical of class VIII bHLH proteins, in both CbbHLHVIII and CnbHLHVIII (Figure 2.C and Supplementary Figure S3 and S4).

**Figure 2.**
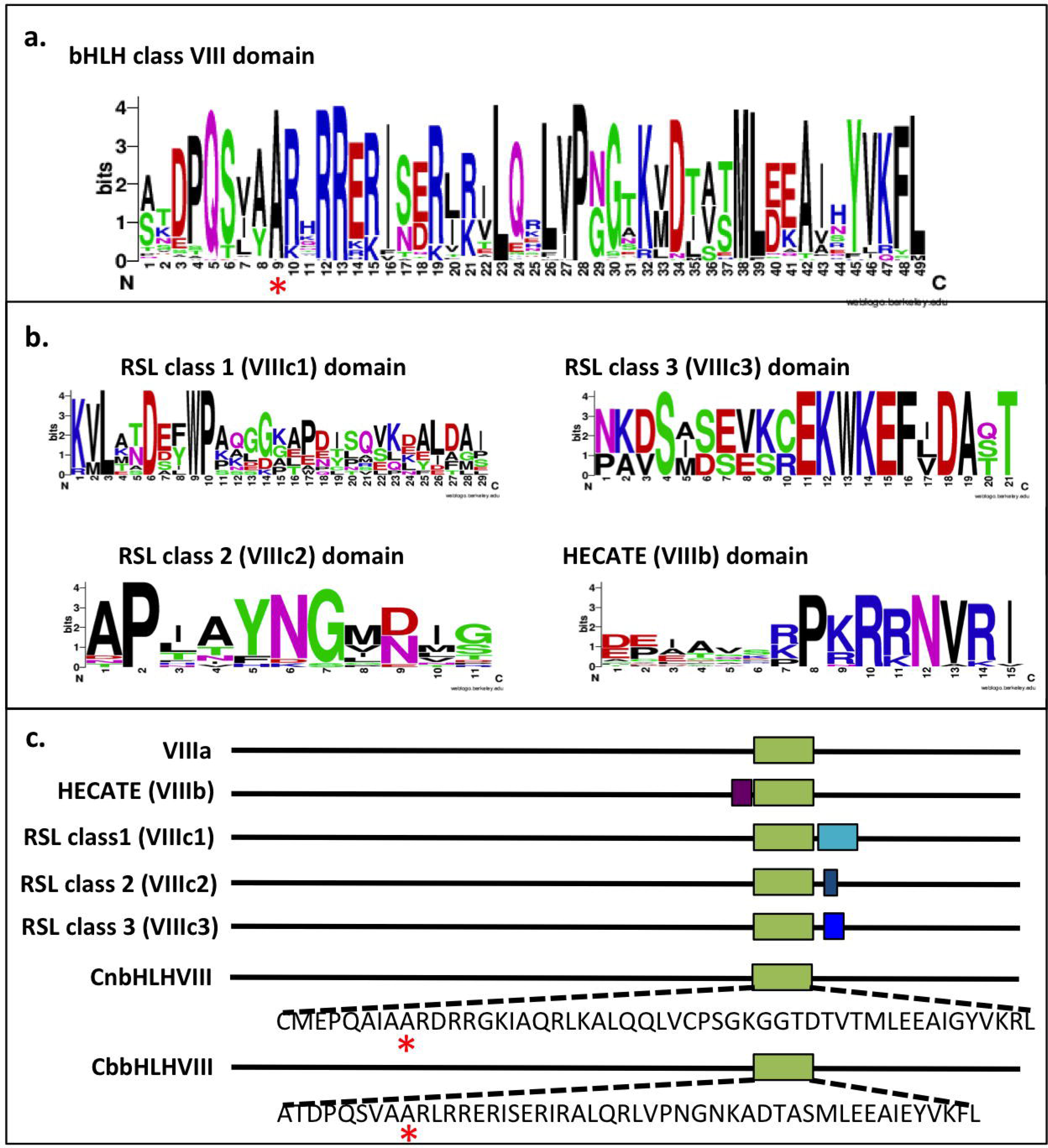
Conserved amino acid motifs of the class VIII bHLH family. **a.** LOGO representation of the amino acid sequence of the atypical bHLH domain of class VIII proteins. The red star marks the position of the conserved Alanine (A^210^ in AtRHD6) specific of the class VIII bHLH proteins. **b.** LOGO representations of the amino acid sequences of the conserved motifs of the class VIII bHLH subclasses. **c.** Position of the conserved class VIII bHLH amino acid domains in the land plant class VIII proteins and CbbHLHVIII and CnbHLHVIII. Class VIII bHLH domain (green box), HECATE domain (purple box), RSL domains (blue boxes). The sequence of the *C. braunii* and *C. nitellarum* bHLH domains are given. Red stars mark the position of the conserved Alanine (A^210^ in AtRHD6).

There are conserved characteristic amino acid motifs in RSL (subclass VIIIc) and HECATE (subclass VIIIb) proteins (Pires & Dolan, 2010a). A conserved motif is located just after the bHLH domain in members of the RSL subfamily (VIIIc) (Figure 2.C and Supplementary Figure S3). The precise sequence of this motif is characteristic of each of the three monophyletic RSL subclasses. KVLATDEFWPAQGGKAPDISQVKDALDAI is found in members of the RSL subclass 1 (VIIIc1), APIAYNGMDIG in members of the RSL subclass 2 (VIIIc2) and NKDSASEVKCEKWKEFIDAQT in members of the RSL subclass 3 (VIIIc3) (Figure 2.B). Similarly, a conserved sequence (DPIAVSRPKRRNVRI) is located just before the N-terminal of the bHLH domain in members of the HECATE (VIIIb) subclass (Figure 2.B and C). None of the conserved amino-acid motifs of the RSL and HECATE proteins are present in the CbbHLHVIII or CnbHLHVIII protein sequences (Figure 2.C and Supplementary Figure S3). This suggests that these land plant specific motifs evolved after the divergence of *C. braunii* and *C. nitellarum* from the last common ancestors with the land plants.

### CbbHLHVIII protein cannot replace the RSL proteins that positively regulate root hair development in *Arabidopsis thaliana*

To assess if Cb*bHLHVIII* controls the development of filamentous rooting cells (rhizoids and root hairs) in land plants, we tested if Cb*bHLHVIII* could substitute for the loss of RSL function in *A. thaliana* mutants. At*rhd6* At*rsl1* double mutants are devoid of RSL class 1 function and do not develop root hairs (Figure 3) (Menand *et al.*, 2007). To test the ability of Cb*bHLHVIII* to restore root hair development, At*rhd6* At*rsl1* double mutants were transformed with a gene construct driving the expression of Cb*bHLHVIII* under the control of the cauliflower mosaic virus 35S constitutive promoter (p35S:Cb*bHLHVIII:T*). We identified three lines that expressed Cb*bHLHVIII* at high levels (Figure 3.A and B). None of these At*rhd6* At*rsl1* p35S:Cb*bHLHVIII:T* plants developed root hairs (Figure 3.C). As a control, At*rhd6* At*rsl1* double mutants were transformed with either a p35S:AtRHD6:T or a p35S:MpRSL1:T gene constructs which overexpressed the A. thaliana and M. polymorpha RSL subclass 1 genes respectively (Figure 3.A and B). Both the Atrhd6 Atrsl1 p*35S*:At*RHD6:T* and the At*rhd6* At*rsl1* p*35S*:Mp*RSL1:T* plants developed root hairs (Figure 3.C). These data indicate that expression of a class VIII protein from *C. braunii* cannot compensate for loss of At*RHD6* and At*RSL1* function. This suggests CbbHLHVIII does not function in rhizoid development and indicates that CbbHLHVIII is functionally different from the RSL subclass 1 proteins AtRHD6 and AtRSL1.

**Figure 3.**
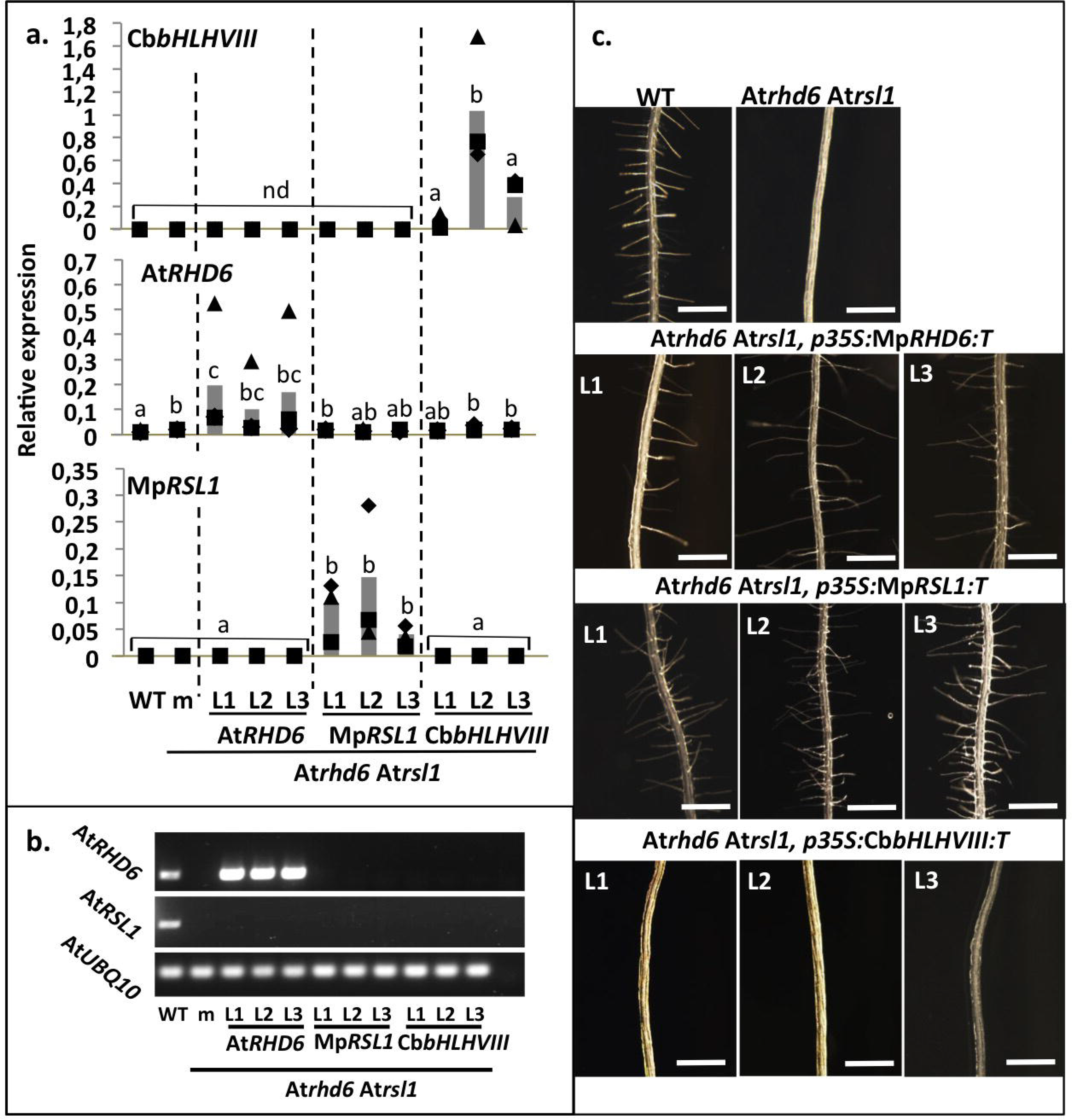
Cb*bHLHVIII* expression does not restore root hair development on root hairless *Arabidopsis thaliana* At*rhd6* At*rsl1* mutants. **a.** Histograms showing the mean steady state levels *(n=3)* of At*RHD6*, Mp*RSL1* and Cb*bHLHVIII* mRNA in *Arabidopsis thaliana* wild type (WT), At*rhd6* At*rsl1* double mutants and At*rhd6* At*rsl1* double mutants transformed with p*35S*:At*RHD6:T* (At*RHD6*), p*35S*:Mp*RSL1:T (*Mp*RSL1)* or p*35S*:Cb*bHLHVIII:T (*Cb*bHLHVIII)*. L1, L2 and L3 are three independently transformed lines for each transgene. Transcripts levels were normalized to At*UBQ10* and At*PDF2* mRNA levels. Each biological replicate is represented by a square, a triangle and a diamond for replicate 1, replicate 2 and replicate 3, respectively. In some lines the expression of Cb*bHLHVIII* was not detected (nd). Different letters refers to statistically different groups (P<0,05, Kruskall-Wallis test). **b.** Analysis of the presence or absence of At*RHD6* and At*RSL1* transcripts by RT-PCR in *Arabidopsis thaliana* wild type (WT), At*rhd6* At*rsl1* double mutants and Atrhd6 Atrsl1 double mutants transformed with p*35S*:At*RHD6:T (*At*RHD6*), p*35S*:Mp*RSL1:T* (Mp*RSL1*) or p*35S*:Cb*bHLHVIII:T (*Cb*bHLHVIII*) constructs. L1, L2 and L3 are three independently transformed lines for each transgene. At*UBQ10* is used as a reference gene. **c.** Root hair phenotypes of Arabidopsis thaliana wild type (WT), At*rhd6* At*rsl1* double mutant and three independent lines (L1, L2, L3) for each of the At*rhd6* At*rsl1* double mutants transformed with p*35S*:At*RHD6:T*, p*35S*:Mp*RSL1:T* or p*35S*:Cb*bHLHVIII:T constructs*. Scale bars: 1 mm.

### CbbHLHVIII protein cannot replace the RSL protein that positively regulates rhizoid development in *Marchantia polymorpha*

To independently determine if the function of CbbHLHVIII could substitute for RSL function in land plants, we tested the ability of CbbHLHVIII to restore rhizoid development in the rhizoidless Mp*rsl1* loss of function mutant of *M. polymorpha* (Figure 4 and 5). We transformed the Mp*rsl1* mutant with a gene construct in which the Cb*bHLHVIII* gene was under the control of the constitutive promoter of *M. polymorpha* EF1*α* (pMp*EF1α*: Cb*bHLHVIII:T*) (Althoff *et al.*, 2014). Five Mp*rsl1* pMp*EF1α*:Cb*bHLHVIII:T* lines with high steady state levels of the *C. braunii* transgene were identified (Figure 4). None of these Mp*rsl1* pMp*EF1α*:Cb*bhHLH:T* plants developed rhizoids (Figure 5). As a control, we transformed the Mp*rsl1* loss of function mutant with a pMp*EF1α*:Mp*RSL1:T* gene construct. Mp*rsl1* plants transformed with pMp*EF1α*:MpR*SL1:T* developed rhizoids indicating that overexpression of the Mp*RSL1* class VIII protein driven by the promoter of *M. polymorpha* EF1*α* compensate for the loss of Mp*RSL1* function (Figure 5). The lack of rhizoid development on Mp*rsl1* pMp*EF1α*:Cb*bHLHVIII:T* plants demonstrates that class VIII bHLH protein from *Chara braunii* (Cb*bHLHVIII*) cannot substitute for Mp*RSL1* function during rhizoid development in Mp*rsl1* mutants. These data suggest that Cb*bHLHVIII* and Mp*RSL1* are functionally different.

**Figure 4.**
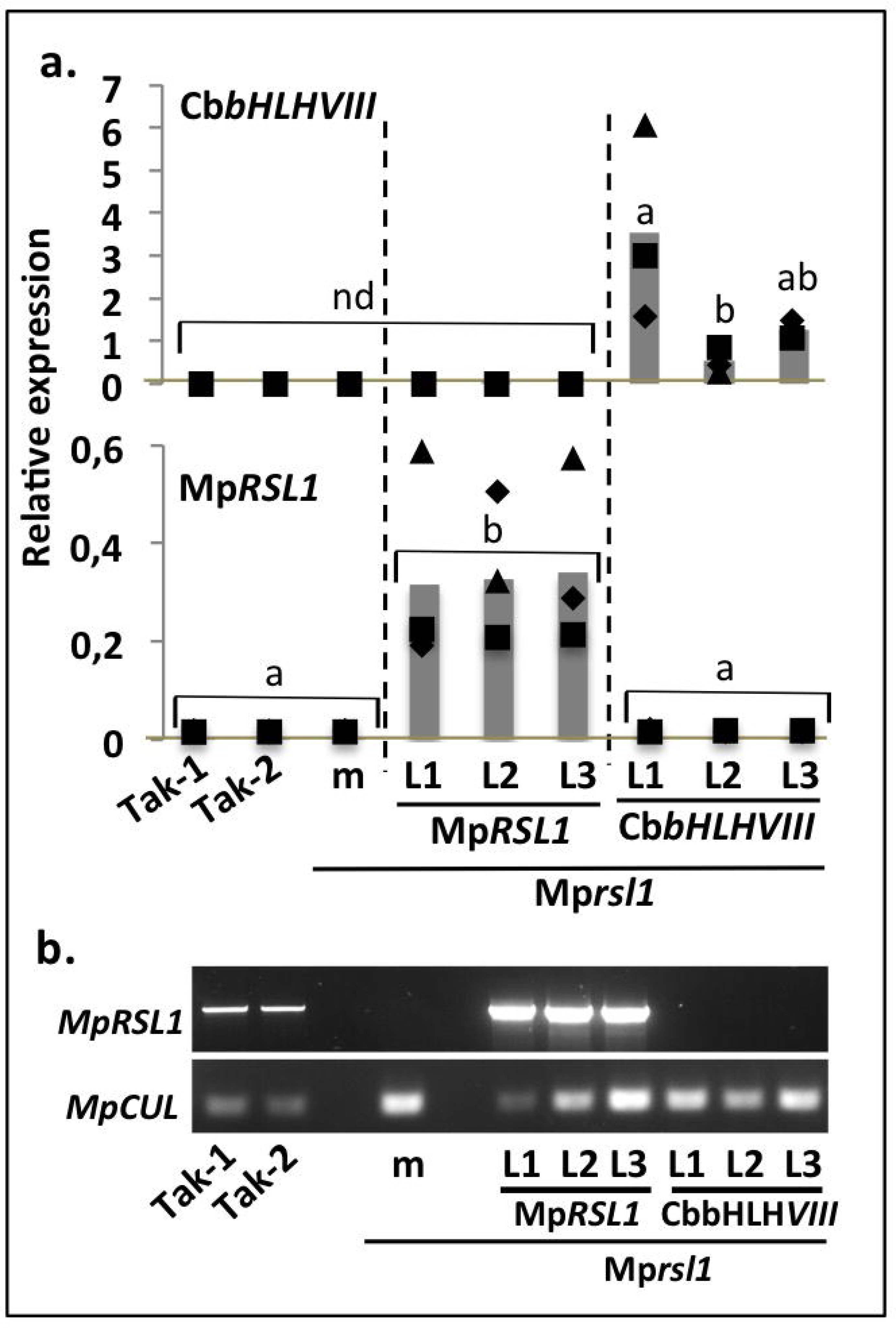
Cb*bHLHVIII* expression in *Marchantia polymorpha* Mp*rsl1* mutants. **a.** Histograms showing the mean steady state levels *(n=3)* of At*RHD6*, Mp*RSL1* and Cb*bHLHVIII* mRNA in *M. polymorpha* wild type male (Tak-1) and female (Tak-2), Mp*rsl1* mutant (m) and Mp*rsl1* mutants transformed with pMp*EF1α*:Mp*RSL1:T* (Mp*RSL1*) or pMp*EF1α*:Cb*bHLHVIII:T* (Cb*bHLHVIII*). For each construct, L1, L2 and L3 are three independently transformed lines. Transcripts levels were normalized to Mp*ACT* and Mp*CUL* mRNA levels. Each biological replicate is represented by a square, a triangle and a diamond for replicate 1, replicate 2 and replicate 3, respectively. Cb*bHLHVIII* was not detected (nd) in some lines. Different letters refers to statistically different groups (P<0,05, Kruskall-Wallis test). **b.** Analysis of the presence or absence of MpRSL1 transcript by RT-PCR in Marchantia polymorpha wild type male (Tak-1) and female (Tak-2), Mp*rsl1* mutant (m) and Mp*rsl1* mutants transformed with pMp*EF1α*:Mp*RSL1:T* (Mp*RSL1*) and pMp*EF1α*:Cb*bHLHVIII:T* (Cb*bHLHVIII*). Mp*CUL* is the reference gene. L1, L2 and L3 are three independently transformed lines.

**Figure 5.**
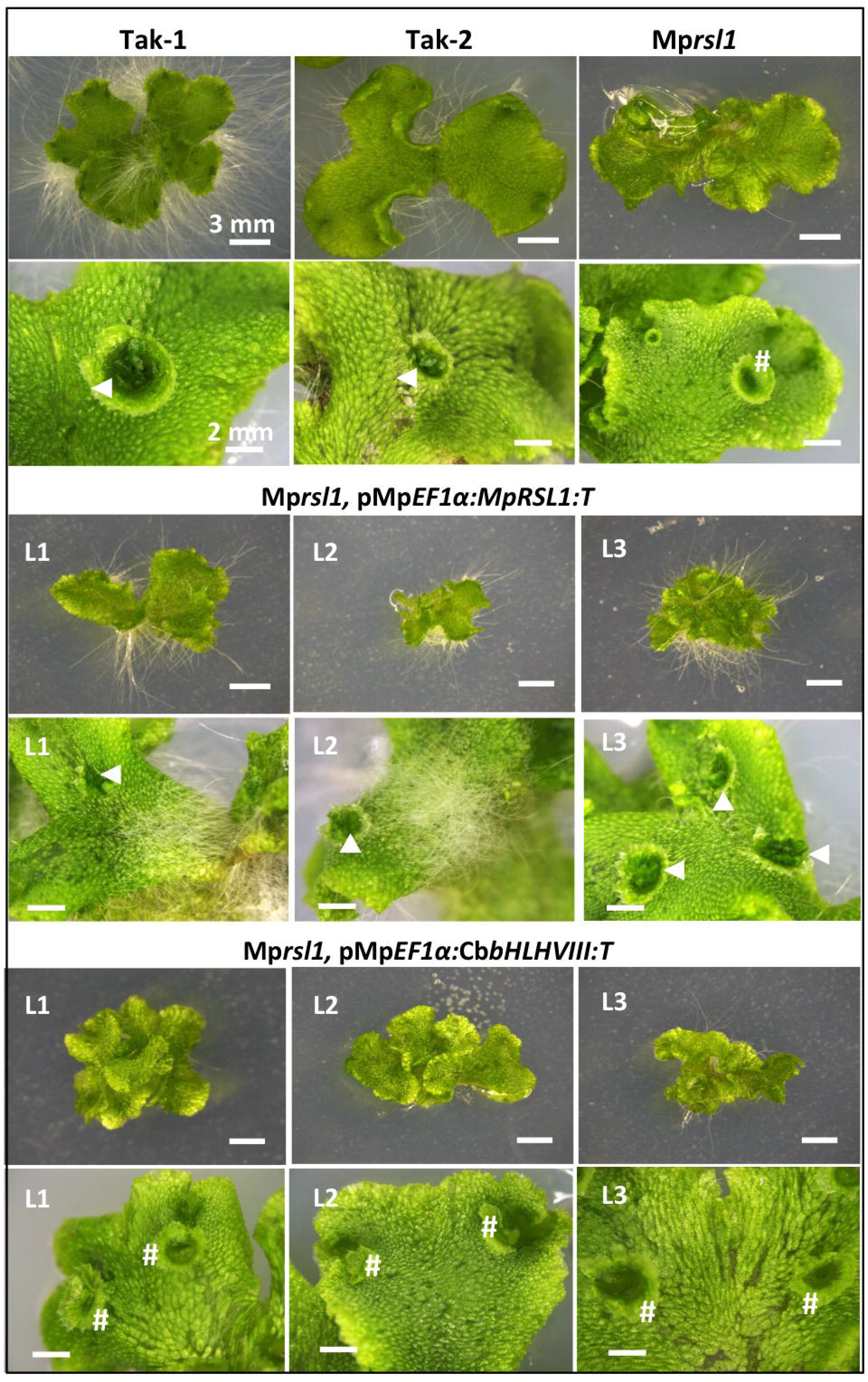
Cb*bHLHVIII expression in M. polymorpha* Mp*rsl1* mutants does not restore rhizoid and gemmae development. Rhizoid and gemma cup phenotypes of wild type male (Tak-1) and female (Tak-2) *M. polymorpha*, Mp*rsl1* mutants and three independent lines (L1, L2, L3) of Mp*rsl1* mutants transformed with pMp*EF1α*:MpR*SL1:T* or pMp*EF1α*:Cb*bHLHVIII:T*. For each genotype the top image represents the rhizoid phenotype of regenerated thalli seven weeks after transformation. Scale bars: 3 mm. The bottom images are the gemma cups of regenerated thalli ten weeks after transformation. White arrows indicate gemma cups full of gemmae. # indicates empty gemma cups. Scale bars: 2 mm.

### Expression of Cb*bHLHVIII* in wild type *Marchantia polymorpha* does not induce supernumerary rhizoid development

To verify that Cb*bHLHVIII* could not promote rhizoid development in *M. polymorpha* using a different experimental approach, we ectopically overexpressed Cb*bHLHVIII* in wild type and compared the phenotypes to the phenotypes of plants that ectopically overexpress the endogenous Mp*RSL1* gene (Figure 6). Ectopic overexpression of Mp*RSL1* using the 35S promoter (p*35S*:Mp*RSL1:T*) induced the development of supernumerary rhizoids in wild type. However, expression of Cb*bHLHVIII* from the 35S promoter (p*35S*:Cb*bHLHVIII:T*) in wild type *M. polymorpha* did not induce supernumerary rhizoid development. This verifies that Cb*bHLHVIII* cannot function during rhizoid development in *M. polymorpha*. This is consistent with the hypothesis that Cb*bHLHVIII* does not control rhizoid development and that CbbHLHVIII and MpRSL1 proteins are functionally different.

**Figure 6.**
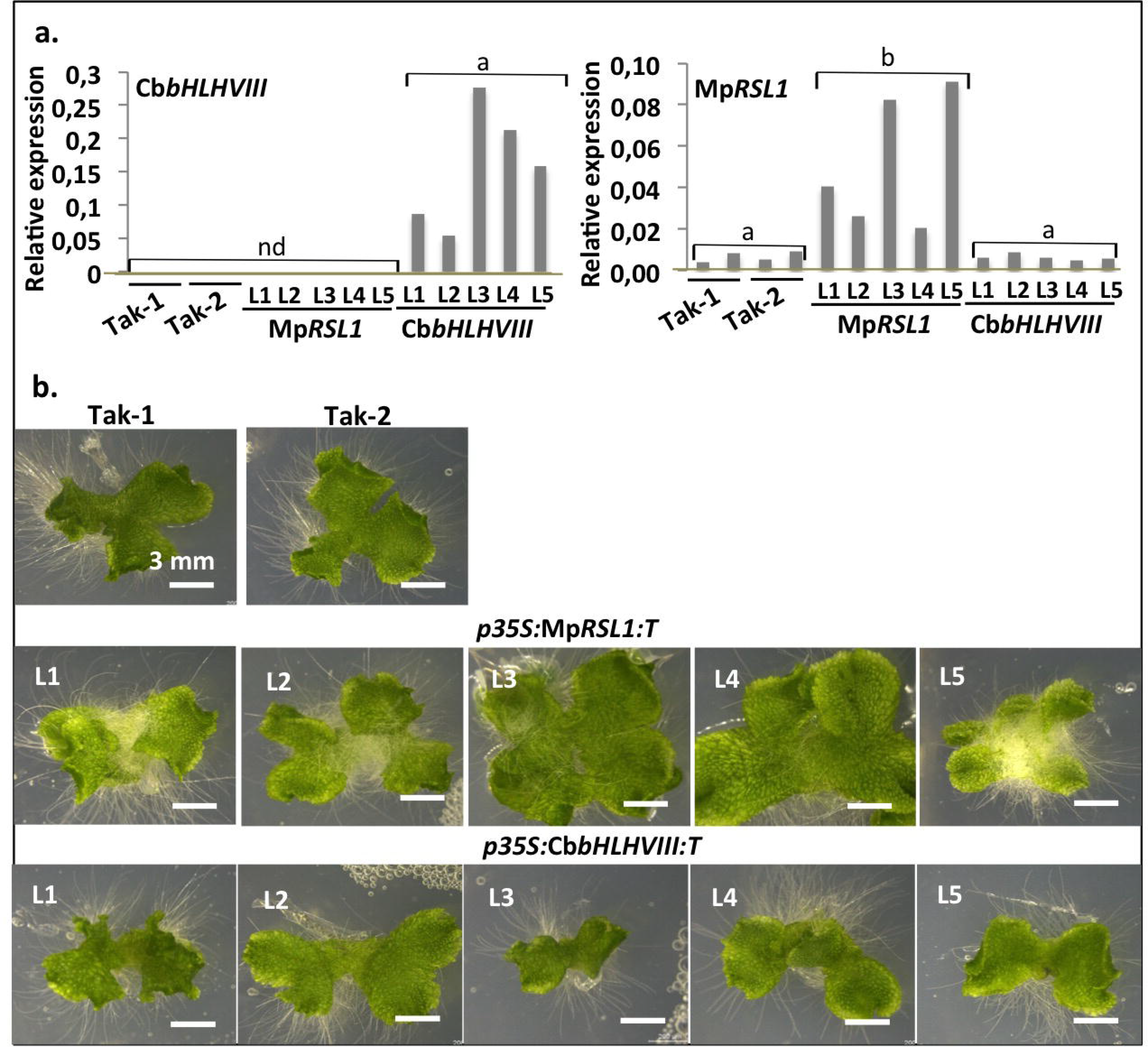
Expression of Cb*bHLHVIII* in *Marchantia polymorpha* does not induce supernumerary rhizoid development. **a.** Histograms showing the steady state level of Mp*RSL1* and Cb*bHLHVIII* mRNA in *M. polymorpha* wild type male (Tak-1) and female (Tak-2) and wild type *M. polymorpha* transformed with p35S:Cb*bHLHVIII:T* (Cb*bHLHVIII*) or p*35S*:Mp*RSL1:T* (Mp*RSL1*). For each construct, five lines (L1, L2, L3, L4 and L5) were independently transformed. Transcripts levels were normalized with the geometrical mean of Mp*ACT* and Mp*CUL* expression levels. In some lines the expression of Cb*bHLHVIII* was not detected (nd). Different letters refers to statistically different groups (P<0,05, Kruskall-Wallis test). **b.** Rhizoid phenotypes of *M. polymorpha* WT male (Tak-1) and female (Tak-2) and *M. polymorpha* wild type transformed with p*35S*:Mp*RSL1:T* or p*35S*:Cb*bHLHVIII:T*. Plants are four weeks old thalli grown from spores that were transformed with transgenes. Scale bars: 3 mm.

### CbbHLHVIII protein cannot replace the RSL protein that positively regulates gemma development in *Marchantia polymorpha*

To independently verify that CbbHLHVIII cannot substitute for MpRSL1 in *M. polymorpha*, we tested the ability of Cb*bHLHVIII* to restore gemma development in Mp*rsl1* mutants (Figure 5). Mp*rsl1* mutants rarely develop gemmae; the gemma cups of Mp*rsl1* mutants are empty while gemmae can fill gemma cups of wild type plants. None of the Mp*rsl1* plants transformed with the pMp*EF1α*:Cb*bHLHVIII:T* gene construct developed gemmae and gemma cups were empty. The control Mp*rsl1* plants transformed with the pMp*EF1α*:Mp*RSL1:T* gene construct developed gemmae and gemma cups were full. Together these data indicate that overexpression of the Cb*bHLHVIII* protein cannot compensate for loss of Mp*RSL1* function during gemma development in Mp*rsl1* mutants. This is consistent with the hypothesis that the function of Cb*bHLHVIII* is different from Mp*RSL1*.

### *Chara braunii* develop rhizoids from multicellular nodes

The green thallus of *C. braunii* comprises several axes, each consisting of alternating nodes and internodes (Figure 7.A) (Smith & Allen, 1955; Pickett-Heaps, 1975; Graham & Wilcox, 2000; Nishiyama *et al.*, 2018). The internodes are composed of a single elongated cell, while the nodes are multi-cellular and are the sites from which branches (determinate structures) and new axes (indeterminate structures) initiate. To define where *C. braunii* rhizoids develop, we grew individual *C. braunii* algae in axenic liquid culture and found that rhizoids developed from the nodal complexes (Figure 7). No rhizoids were present on nodes 1 and 2 where node 1 is the node nearest the apex (Figure 7.B). After initiation from a nodal cell, the rhizoid elongates by tip growth (Supplementary Figure S5) and cell division. Growing rhizoids can branch (Supplementary Figure S5). Rhizoids of all developmental stages – initiating rhizoids, elongating rhizoids, multicellular rhizoids and branched rhizoids – are present on node 3 and older nodes (Figure 7.B). These observations indicate that, in our growth condition, rhizoids develop from the nodal complexes of the thallus. Nodal initiation of rhizoids begins from the nodes in 3^rd^ position from the apical meristem and initiation continues in older nodes (Figure 7.B and C).

**Figure 7.**
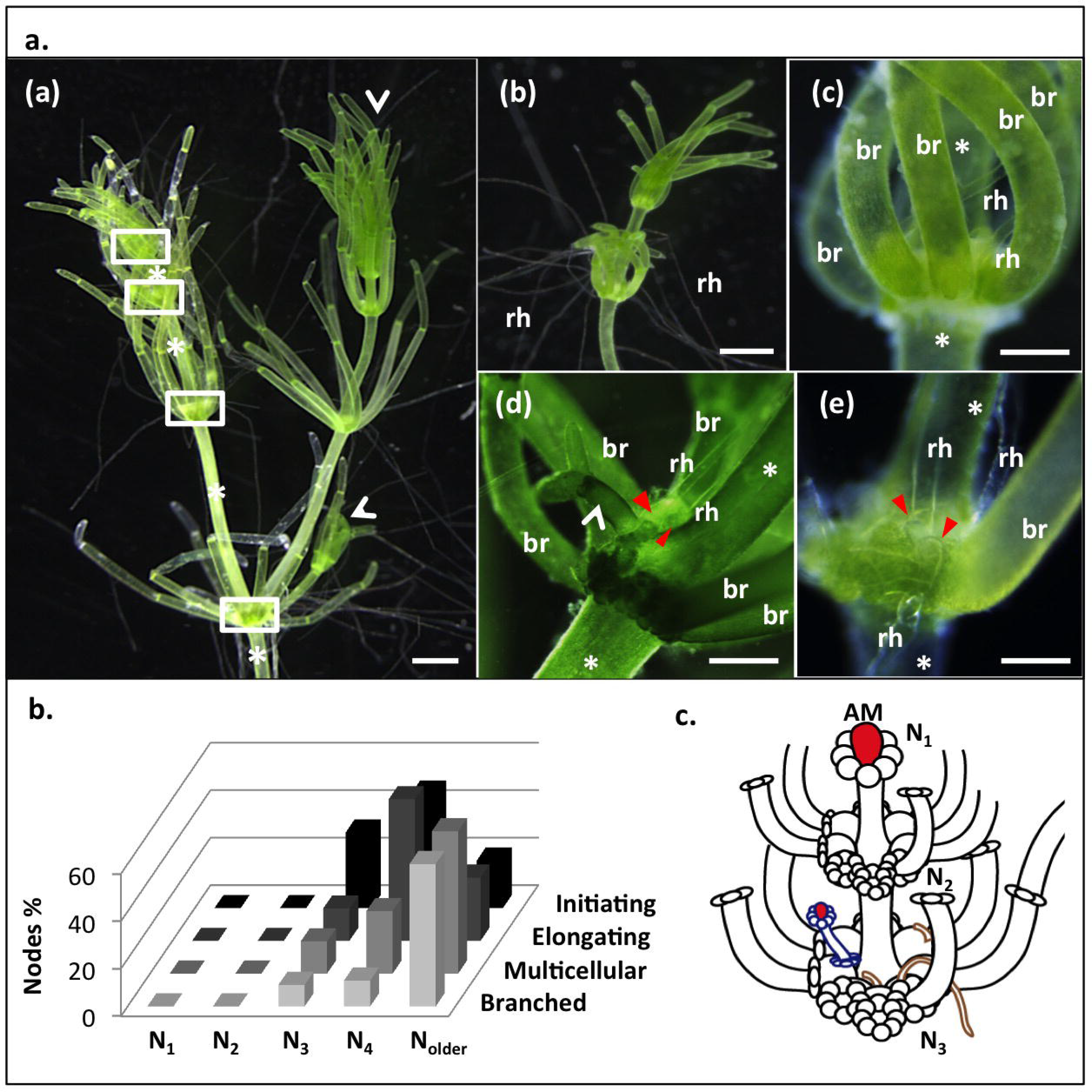
Rhizoids develop on *Chara braunii* nodal complexes. **a.** *C. braunii* thallus. (a) Nodal complexes on the main axis of the thallus are framed in white. A star indicates the unicellular internodes of the main axis of the thallus. White arrowheads indicate the new lateral axes that developed from nodal complexes. (b) *C. braunii* rhizoids (rh) growing from a nodal complexes. (c),(d) and (e) White stars mark the internodal cells on both sides of the multicellular nodes. Branches (br), new axes (white arrowheads) and rhizoids (rh) develop from the nodal complexes. Red arrowheads indicate where each rhizoid is attached to the nodal complex. Scale bars: 1 mm (a) and (b), 250 μm (c), (d) and (e). Branches were removed in (d) and (e) to reveal the site where rhizoids and branches attach to the node. **b.** Percentages of nodes of different developmental stages bearing rhizoids at different developmental stage (Initiating, elongating, multicellular, branched). N_1_ are the first nodes below the apical meristem, N_2_ are the second, N_3_ are the third, N_4_ are the fourth and N_older_ are the fifth and older. **c.** Schematic of a *C. braunii* axis showing branches attached to the nodes, rhizoids (in brown), new axes (in blue) and apical meristems (in red).

### Cb*bHLHVIII* is expressed at the apex and gametangia of *Chara braunii*

The demonstration that expression of Cb*bHLHVIII* does not restore rhizoid development in At*rhd6* At*rsl1* double mutants or At*rhd6* Ar*tsl1* double mutants suggests that this protein is not involved in rhizoid development in *C. braunii*. In land plants, RSL Class 1 encoding genes (subclass VIIIc) are expressed in filamentous rooting cells – rhizoid cells and root hair cells – and the cells from which rhizoid cells and root hair cells develop (Menand *et al.*, 2007; Jang *et al.*, 2011; Kim & Dolan, 2016; Proust *et al.*, 2016; Kim *et al.*, 2017). To determine where Cb*bHLHVIII* is expressed during *C. braunii* development, we measured the steady state levels of Cb*bHLHVIII* mRNA using quantitative RT-PCR (Figure 8). In the first experiment, RNA was isolated from thallus, rhizoids, gametangiophores (antheridia and archegonia) and zygotes (Figure 7 and Supplementary Figure S6). The highest steady state levels of Cb*bHLHVIII* mRNA were detected in the thallus and gametangiophores (Figure 8.A). Low steady state levels of Cb*bHLHVIII* mRNA were detected in rhizoids and zygotes. These results were consistent with the levels of expression of Cb*bHLHVIII* (Cb_Transcript_119934) detected in the thallus and rhizoid transcriptomes (Supplementary Table 1). To identify where in the thallus Cb*bHLHVIII* mRNA accumulated, we isolated RNA from different thallus structures (Figure 8.B). Steady state levels of Cb*bHLHVIII* mRNA were highest at the apex (apical meristem and node 1) and levels were progressively lower in nodes 2 and nodes 3. Relatively high steady state levels of Cb*bHLHVIII* mRNA were observed in gametangia-bearing branches, while Cb*bHLHVIII* mRNA levels were lower in branches without gametangia. This suggests that Cb*bHLHVIII* mRNA accumulates in apices and gametangia. These data are consistent with Cb*bHLHVIII* being expressed in the morphogenetic centres of *C. braunii*, the apex, node 1 and gametangia. The low steady state levels of Cb*bHLHVIII* mRNA in rhizoids and in nodes bearing initiating rhizoids (node 3 and older) suggest that Cb*bHLHVIII* expression is not involved in rhizoid development. Taken together these data indicate that CbbHLHVIIII is expressed in the *C. braunii* morphogenetic centres but suggest that this transcription factor not involved in rhizoid development.

**Figure 8.**
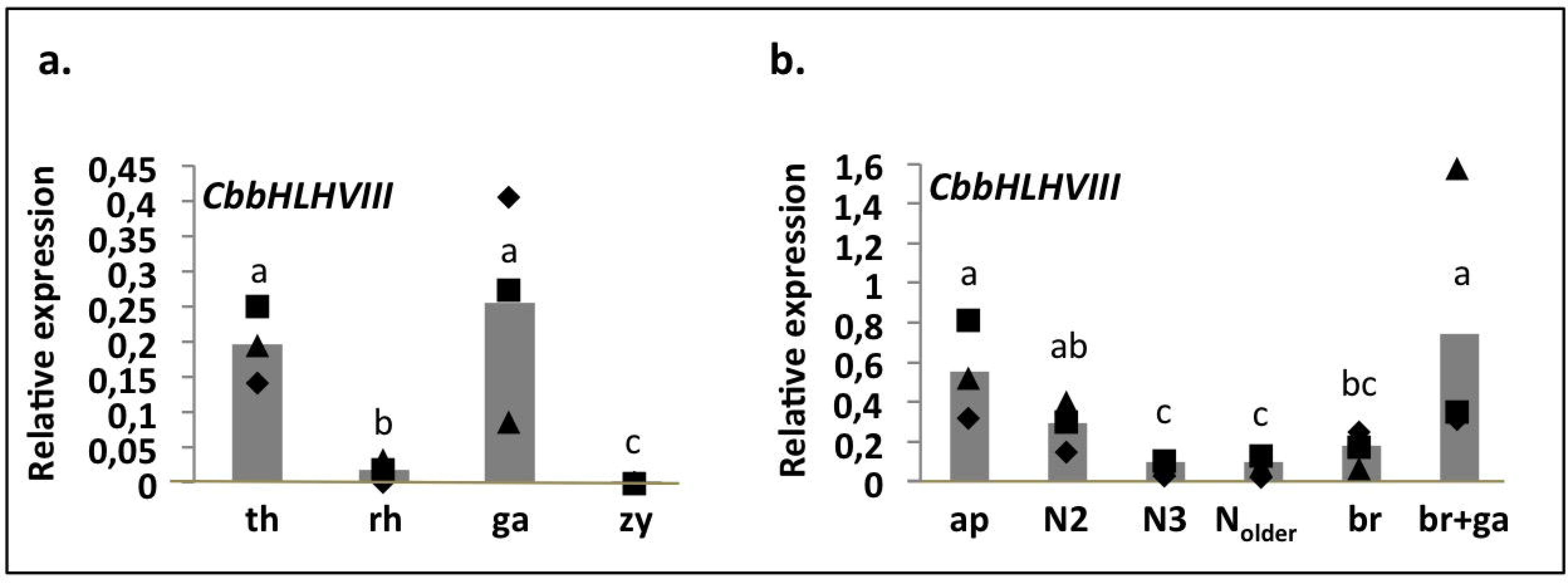
Cb*bHLHVIII* mRNA is expressed in morphogenetic centres. **a.** Histograms showing the mean steady state levels (n=3) of Cb*bHLHVIII* mRNA in *C. braunii* thallus (th), rhizoids (rh), gametangia (ga) and zygotes (zy). Transcripts levels were normalized with Cb*ELF5a* expression. Each biological replicate is represented by a square, a triangle and a diamond for replicate 1, replicate 2 and replicate 3, respectively. Different letters refer to statistically different groups (P<0,05, Kruskall-Wallis test). **b.** Histogram showing the mean steady state level of Cb*bHLHVIII* mRNA in different regions of the *C. braunii* thallus: the apex (apical cells and first nodal complex; ap), nodes 2 (N2), nodes 3 (N3), older nodes (N*older*), branches (br) and branches bearing gametangia (br+ga). Transcript levels were normalized to Cb*ELF5a* mRNA levels. Each biological replicate is represented by a square, a triangle and a diamond for replicate 1, replicate 2 and replicate 3, respectively. Different letters refers to statistically different groups (P<0,05, Kruskall-Wallis test).

## DISCUSSION

Charophycean algae develop complex bodies from morphogenetic centres located at the apices of their axes, like land plants (Smith & Allen, 1955; Pickett-Heaps, 1975; Graham & Wilcox, 2000). These axes develop a diversity of cell types including tip-growing cells called rhizoids that are involved in nutrient uptake (Box, 1986, 1987; Andrews, 1987; Vermeer *et al.*, 2003; Wuestenberg *et al.*, 2011) and anchorage (Graham & Wilcox, 2000). Tip-growing cells involved in nutrient uptake and anchorage also evolved among the land plants (Jones & Dolan, 2012; Bonnot *et al.*, 2017): the tip-growing rhizoids of bryophytes and root hairs of euphyllophytes. No other streptophyte algae form morphogenetic centres or develop the same level of cellular complexity that is characteristic of the charophycean algae (Pickett-Heaps, 1975; Graham & Wilcox, 2000). The distribution of morphological traits on the most recent streptophyte phylogenies suggests that increased body plan complexity evolved independently in the charophycean lineage and in the lineage leading to the land plants (embryophytes) (Wodniok *et al.*, 2011; Wickett *et al.*, 2014; Nishiyama *et al.*, 2018). If rhizoid evolved independently in the land plants and charophycean lineages, we might predict that different mechanisms regulating rhizoid differentiation evolved in the two lineages.

Two broad conclusions can be made from our results. First, we conclude that different mechanisms control the initiation of rhizoid development in Charophycean algae and land plants. Second, we conclude that the acquisition of the rhizoid development function by class VIII bHLH evolved in the land plant lineage after the divergence of extant streptophyte algae and the land plants from their last common ancestor.

Class VIII bHLH proteins are required for the formation of tip-growing rooting cells in in land plants (Masucci & Schiefelbein, 1994; Menand *et al.*, 2007; Jang *et al.*, 2011; Kim & Dolan, 2016; Proust *et al.*, 2016; Kim *et al.*, 2017). Here, we identified a class VIII bHLH protein (Cb*bHLHVIII*) gene from the charophycean alga *C. braunii* that is sister to land plant RSL genes on protein trees. Cb*bHLHVIII* transcripts accumulate at high levels in parts of the plant undergoing morphogenesis, including the apex, the nodes and the gametangia. However, the expression of Cb*bHLHVIII* was very low in rhizoids and the transcript was hardly detectable in regions of the plant from which rhizoids develop. This suggests that Cb*bHLHVIII* does not function in rhizoid development in *C. braunii*. This conclusion is supported by the observation that a single class VIII bHLH protein is also present in the genome of *C. nitellarum*, which does not develop rhizoids (Pickett-Heaps, 1975; Graham & Wilcox, 2000). These data support the hypothesis that class VIIIc bHLH proteins carry out different functions in streptophyte algae and land plants.

Further evidence supporting the hypothesis that Cb*bHLHVIII* does not control rhizoid development is its inability to promote rhizoid development when expressed in land plants. CbbHLHVIII does not substitute for loss of RSL class 1 function in land plant rsl loss of function mutants. This indicates that Cb*bHLHVIII* is functionally different from land plant RSL class 1 genes. This finding is further supported by the inability of Cb*bHLHVIII* expression to restore gemma development in *rsl* loss of function mutants of *M. polymorpha*. These results suggest that RSL class 1 proteins acquired the ability to promote the development of surface structures from single epidermal cells, including rhizoids, after *C. braunii* and land plants last shared a common ancestor. If correct, it would suggest that class VIII proteins evolved novel functions during or after the transition to land. It is possible that these novel functions are conferred by the RSL motifs that are conserved among all land plant RSL proteins but are not present in the streptophyte alga (*C. braunii*, *C. nitellarum*) RSL proteins.

An alternative, though less parsimonious, hypothesis is that the ancestral function of class VIII bHLH transcription factors was to control the development of structures from single cells as it is in land plants, but that this function was lost in both the *C. braunii* and *C. nitellarum* lineages. Ultimately, discovering the function of class VIII bHLH proteins in *Chara* species and *Coleochaete* species requires the functional characterisation of their class VIII bHLH genes. While such characterisation is not currently possible in these algae it would demonstrate the ancestral function of these proteins in the last common ancestor using the comparative approach. The function of RSL class 1 genes in land plants is to control the development of structures that develop from single surface cells. If class VIII bHLH transcription factors controlled the development of a similar process in streptophyte algae, it would suggest that this function was ancestral and likely acted in the last common ancestor of these organism and land plants. However, the current data suggest that function of family VIII bHLH transcription factors in the development of streptophyte algae is entirely different from their function in land plants.

Taken together, the data support a model in which there was a single class VIII bHLH protein in the genome of a streptophyte algal ancestor of the land plants. This gene did not function in rhizoid development. Neofunctionalization occurred in the lineage leading to the land plants and was followed by multiple rounds of gene duplication (Pires & Dolan, 2010a,b; Catarino *et al.*, 2016). These duplicated genes control the development of diverse structures, such as specialised cells and organs derived from single epidermal cells including rhizoids and root hairs (*RSL* genes) (Masucci & Schiefelbein, 1994; Menand *et al.*, 2007; Jang *et al.*, 2011; Kim & Dolan, 2016; Proust *et al.*, 2016; Kim *et al.*, 2017) as well as complex organ development (*HECATE* genes) (Liljegren *et al.*, 2004; Gremski *et al.*, 2007; Kay *et al.*, 2013). This is consistent with the hypothesis that the increase in morphological diversity that accompanied the colonisation of the land resulted from gene duplication and neofunctionalisation of ancestral regulatory genes.

## ACKNOWLEGMENTS

This work was supported by a European Research Council Advanced award (EVO500 project no. 25028) and a Marie SklodowskaLJCurie award (PLANTORIGINS project no. 238640) to L.D. A.J.H. was supported by a Biotechnology and Biological Sciences Research Council Doctoral Training Partnership Scholarship (Grant BB/J014427/1). We are grateful to Pr D. Domozych (Skidmore College, NY, USA) for providing *C. nitellarum* and, to Pr H. Sakayama and Pr T. Nishiyama for providing *C. braunii*.

## AUTHOR CONTRIBUTIONS

L.D. and C.B. designed the research and wrote the paper. C.B. and H.B. produced the RNA samples for *C. braunii* transcriptome RNAseq. A.J.H. and S.K. performed *C. braunii* transcriptome assembly. H.B. built the vectors backbones. C.B. and C.C. performed *M. polymorpha* transformations. C.B. performed all other experiments and analysed all the data.

---

**Supplementary Table 1.** Expression of Cb_Transcript_119934 (Cb*bHLHVIII*) in the transcriptome.

| Libraries | Expression level (TPM) |  |  |  |  |  |  |  |
| --- | --- | --- | --- | --- | --- | --- | --- | --- |
|  | Th1 | Th 2 | Rh1 | Rh2 | Rh3 | Rh4 | WP1 | WP2 |
| Cb_Transcript_119934 | 0.95 | 2.10 | 0.04 | 0.00 | 0.15 | 0.01 | 0.39 | 1.51 |

Expression level in Transcript Per Million (TPM) of the *CbbHLHVIII* transcript (Cb\_Transcript\_119934) in the different libraries used for the transcriptome (two green thallus (Th) libraries, four rhizoids libraries (Rh) and two whole plants libraries (WP)).

**Supplementary Data 1. Transcript, CDS and amino acid sequences of Cb*bHLHVIII* and Cn*bHLHVIII*.**

**Supplementary Data 2. Amino acid alignment of the Archaeplastida bHLH transcription factors.**

**Supplementary Data 3. Trimmed amino acid alignment used for the phylogenetic analysis of the Archaeplastida bHLH transcription factors.**

**Supplementary Data 4. Trimmed amino acid alignment used for the phylogenetic analysis of the class VIII bHLH transcription factors.**

**Supplementary Figure S1. Unrooted maximum-likelihood tree of Archeplastida bHLH transcription factors.**

Cb*bHLHVIII* and Cn*bHLHVIII* in the class VIII bHLH family are highlighted respectively with a red and a blue triangle. The approximate likelihood ratio test (aLRT) support values are included at nodes. The tree includes sequences from the land plants *Arabidopsis thaliana* (At), *Oryza sativa* (Os), *Selaginella moelendorfii* (Sm), *Phycomitrella patens* (Pp) and *Marchantia polymorpha* (Mp); the streptophyte algae *C. braunii*, *C. nitellarum* and *Klebsormidum flaccidum* (Kf), the chlorophyte algae *Volvox carteri* (Vc), *Chlamydomonas reinhardtii* (Cr), *Chlorella variabilis* (Cv), *Ostreococcus tauri* (Ot); the rodophyte alga *Cyanidioschyzon merolae* (Cm).

**Supplementary Figure S2. Maximum-likelihood tree of the class VIII bHLH transcription factors.**

The class VIII bHLH tree is rooted with the bHLH class XIII and XIV families. It includes sequences from the land plants *Arabidopsis thaliana* (At), *Oryza sativa* (Os), *Selaginella moelendorfii* (Sm), *Phycomitrella patens* (Pp) and *Marchantia polymorpha* (Mp); streptophyte algae *C. braunii*, *C. nitellarum* and *Klebsormidum flaccidum* (Kf); the chlorophyte algae *Volvox carteri* (Vc), *Chlamydomonas reinhardtii* (Cr), *Chlorella variabilis* (Cv) and *Ostreococcus tauri* (Ot); and the rodophyte alga *Cyanidioschyzon merolae* (Cm). The positions of Cb*bHLHVIII* and Cn*bHLHVIII* in the class VIII bHLH family are indicated respectively with a blue and a red triangle. The approximate likelihood ratio test (aLRT) support values are included at nodes.

**Supplementary Figure S3. Conserved domains of the bHLH family VIII proteins.**

Location and amino acid sequences of the conserved domains in the class VIII bHLH proteins. The conserved domains are framed in black (bHLH domain), purple (HECATE domain), light blue (RSL class 1/VIIIc1), grey blue (RSL class 2/VIIIc2) and navy blue (RSL class 3/VIIIc3). The red star marks the position of the conserved Alanine (A^210^ in AtRHD6).

**Supplementary Figure S4. LOGO representation of the conserved amino acid sequences of the bHLH domain of class X, XIII, XIV and XV proteins.**

**Supplementary Figure S5. *Rhizoid development on C. braunii*.**

Rhizoid developmental stages. (a) Initiating rhizoid indicated with a red arrowhead. (b) Elongating rhizoid. (c) Multicellular rhizoid with cell walls indicated with an arrowhead. (d) Branched rhizoid. Scale bars: 250 μm.

**Supplementary Figure S6. *C. braunii* gametangia and zygotes.**

(a) and (b) *C. braunii* gametangia (archegonia (ar) and anteridia (an)) develop on the branches (br). (c) and (d) Zygote (zy) retained on the branches (br). Scale bars: 500 μm (a) and (d) and 250 μm (b) and (c).

